# The histone deacetylase HDA15 interacts with MAC3A and MAC3B to regulate intron retention of ABA-responsive genes

**DOI:** 10.1101/2020.11.17.386672

**Authors:** Yi-Tsung Tu, Chia-Yang Chen, Yi-Sui Huang, Ming-Ren Yen, Jo-Wei Allison Hsieh, Pao-Yang Chen, Keqiang Wu

## Abstract

Histone deacetylases (HDAs) play an important role in transcriptional regulation involved in multiple biological processes. In this study, we investigate the function of HDA15 in abscisic acid (ABA) responses. Immunopurification coupled with mass spectrometry-based proteomics was used to identify the HDA15 interacting proteins. We found that HDA15 can interact with the core subunits of MOS4-Associated Complex (MAC), MAC3A and MAC3B. In addition, ABA enhances the interaction of HDA15 with MAC3B. *hda15* and *mac3a/mac3b* mutants are ABA-insensitive in seed germination and hyposensitive to salinity. RNA sequencing (RNA-seq) analysis demonstrate that HDA15 and MAC3A/MAC3B not only affect the expression of ABA-related genes, but also regulate ABA-responsive intron retention (IR). Furthermore, HDA15 and MAC3A/MAC3B reduce the histone acetylation level of the genomic regions near ABA-responsive IRs. Our studies uncovered the role of histone deacetylation in ABA-mediated splicing regulation and identified that HDA15-MAC3A/MAC3B acts as an important regulation module to mediate splicing of introns in ABA responses.

**One Sentence Summary:** HDA15 and MAC3A/MAC3B coregulate intron retention and reduce the histone acetylation level of the genomic regions near ABA-responsive retained introns.

## Introduction

Post-translational modifications of histones including methylation, acetylation, phosphorylation, ubiquitination, and SUMOylation play important roles in modulating many essential biological processes such as biotic stress, abiotic stress and flowering in plants (Chen and Tian, 2007; Kim et al., 2015; Ueda and Seki, 2020). Histone acetylation catalyzed by histone acetyltransferases (HATs) is essential for gene activation through relaxing chromatin structure (Fisher and Franklin, 2011). Histone deacetylases (HDAs) can remove the acetyl groups from histones, resulting in chromatin compaction and repression of gene expression. Most of the studies are focused on H3 and H4 acetylation since H3 and H4 are more preferable substrates for HATs and HDAs than H2A and H2B. At least 4 lysine residues (K9, K14, K18 and K23) on H3 and 5 lysine residues (K5, K8, K12, K16 and K20) on H4 can be acetylated and deacetylated by HATs and HDAs (Chen and Tian, 2007). In Arabidopsis, HDAs are grouped into three families: the Reduced Potassium Dependency 3 (RPD3)/HDA1 superfamily, Silent Information Regulator 2 (SIR2) family and HD2 family (Pandey et al., 2002; Liu et al., 2014).

HDA15, a RPD3/HDA1-type histone deacetylase, has been reported to mediate different biological processes including photomorphogenesis, ABA response, and stress tolerance. HDA15 is associated with PIF3 and is involved in the regulation of chlorophyll biosynthesis and photosynthesis by reducing histone H4 acetylation levels in the dark (Liu et al., 2013). HDA15 also interacts with PIF1 to regulate the transcription of the light-responsive genes involved in multiple hormonal signaling pathways and cellular processes in germinating seeds in the dark (Gu et al., 2017). Furthermore, HDA15 acts cooperatively with the NC-YC transcriptional factors to regulates light control of hypocotyl elongation by co-regulating the transcription of the light-responsive genes (Tang et al., 2017). In addition, HDA15 also interacts with HY5 involved in repressing hypocotyl cell elongation during photomorphogenesis(Zhao et al., 2019). More recent studies indicate that HDA15 is involved in abiotic stress responses. HDA15 and HFR1 cooperatively regulate several warm temperature marker genes such as *HSP20* (*HEAT SHOCK PROTEIN 20*), *IAA19* (*INDOLE-3-ACETIC ACID INDUCIBLE 19*), and *IAA29* in plant response to elevated ambient temperature (Shen et al., 2019). Furthermore, MYB96 form a complex with HDA15 to repress the expression of *RHO GTPASE OF PLANTS* (*ROP*) in ABA and drought stress responses (Lee and Seo, 2019). In addition, HDA15 can be phosphorylated in vivo and HDA15 phosphorylation results in the loss of enzymatic activity and functions (Chen et al., 2020).

MAC, a highly conserved complex in eukaryotes, is the orthologous complex of the NINETEEN COMPLEX (NTC) or Prp19 complex (Prp19C) in yeast (Palma et al., 2007). The NTC/Prp19C function is essential for the splicing reaction by regulating the progression of spliceosome rearrangement (Chan et al., 2003; Hogg et al., 2010). The MAC complex consists of over 20 proteins in Arabidopsis, many of them are involved in alternative splicing (AS) (Monaghan et al., 2009; Monaghan et al., 2010; Zhang et al., 2014) and control miRNA levels through modulating pri-miRNA transcription, processing, and stability (Jia et al., 2017; Li et al., 2018). The subunits of the MAC complex, MOS4, CELL DIVISION CYCLE5 (CDC5), PRL1 (PLEIOTROPIC REGULATORY LOCUS 1), MAC3A/3B, and MAC5A/5B/5C are required for plant immunity, since their loss-of-function mutations result in plants exhibiting enhanced susceptibility to pathogen infection (Palma et al., 2007; Monaghan et al., 2009; Monaghan et al., 2010). More recently, it has been found that MAC3A/MAC3B interacts with three splicing factors, SKI-INTERACTING PROTEIN (SKIP), PLEIOTROPIC REGULATORY LOCUS 1 (PRL1), and SPLICEOSOMAL TIMEKEEPER LOCUS1 (STIPL1), to mediate the splicing of pre-mRNAs encoded by circadian clock- and abiotic stress response-related genes (Li et al., 2019).

The involvement of histone modifications in splicing regulation has been reported in yeasts and animals (Pajoro et al., 2017). Two models have been proposed to explain how histone modifications regulate alternative splicing (Rahhal and Seto, 2019). In the kinetic coupling model, histone deacetylation results in the more condensed chromatin, which would slow down the elongation rate. Therefore, the splicing factors are recruited to the weak splicing sites, causing exon inclusion. In contrast, acetylated histones lead to a looser chromatin and a faster elongation rate. As a result, the splicing factors are recruited to the strong splicing sites, leading to exon skipping (Rahhal and Seto, 2019). In chromatin-adaptor model, histone marks recruit different splicing regulators by chromatin-binding protein that reads the histone marks. For example, the heterochromatin 1 protein HP1*r* recognizes H3K9me3 marks to reduce transcriptional elongation rate and include exons in human (Saint-André et al., 2011). However, the role of histone modifications in splicing regulation in plants remains elusive.

In this study, we found that HDA15 interacts with the core subunits of the MAC complex, MAC3A/MAC3B. The loss of function mutants of *HDA15* and *MAC3A/MAC3B* are ABA-insensitive in seed germination, hyposensitive to salinity, and high transpiration rates. Furthermore, transcriptome analyses demonstrate that HDA15 and MAC3A/MAC3B are involved in ABA-mediated splicing regulation, especially IR.

## Results

### Identification of HDA15 interacting proteins

A previous study has shown that HDA15 interacts with the transcription factor MYB96 to regulate the gene expression involved in ABA responses (Lee and Seo, 2019). To further investigate the function of HDA15 in ABA responses, we performed immunopurification coupled with mass spectrometry-based proteomics to identify the HDA15 interacting proteins. 10-day-old Col-0 and *hda15-1* plants treated with or without (mock) ABA were used for immunopurification using an anti-HDA15 antibody. In total, 209 and 435 HDA15-interacting proteins were identified in mock and ABA treatment, respectively (Supplemental Data Set 1). Gene Ontology (GO) analyses indicate that many HDA15-interacting proteins under both ABA-treated and mock conditions are involved in the transcription regulation of the biological processes (Supplemental .Fig. 1A). Moreover, the GO terms, “response to salt stress” and “response to abscisic acid”, are highly enriched in ABA treatment (Supplemental Fig. 1B). The correlation of the interacting proteins was analyzed by STRING (Szklarczyk et al., 2015) and visualized in Cytoscape (Shannon et al., 2003). A large number of HDA15 interacting proteins can be grouped into the transcription factors, chromatin regulators and ribosomal proteins in both mock and ABA-treated conditions (Fig. 1A, B, Supplemental Data Set 1).

**Fig. 1.**
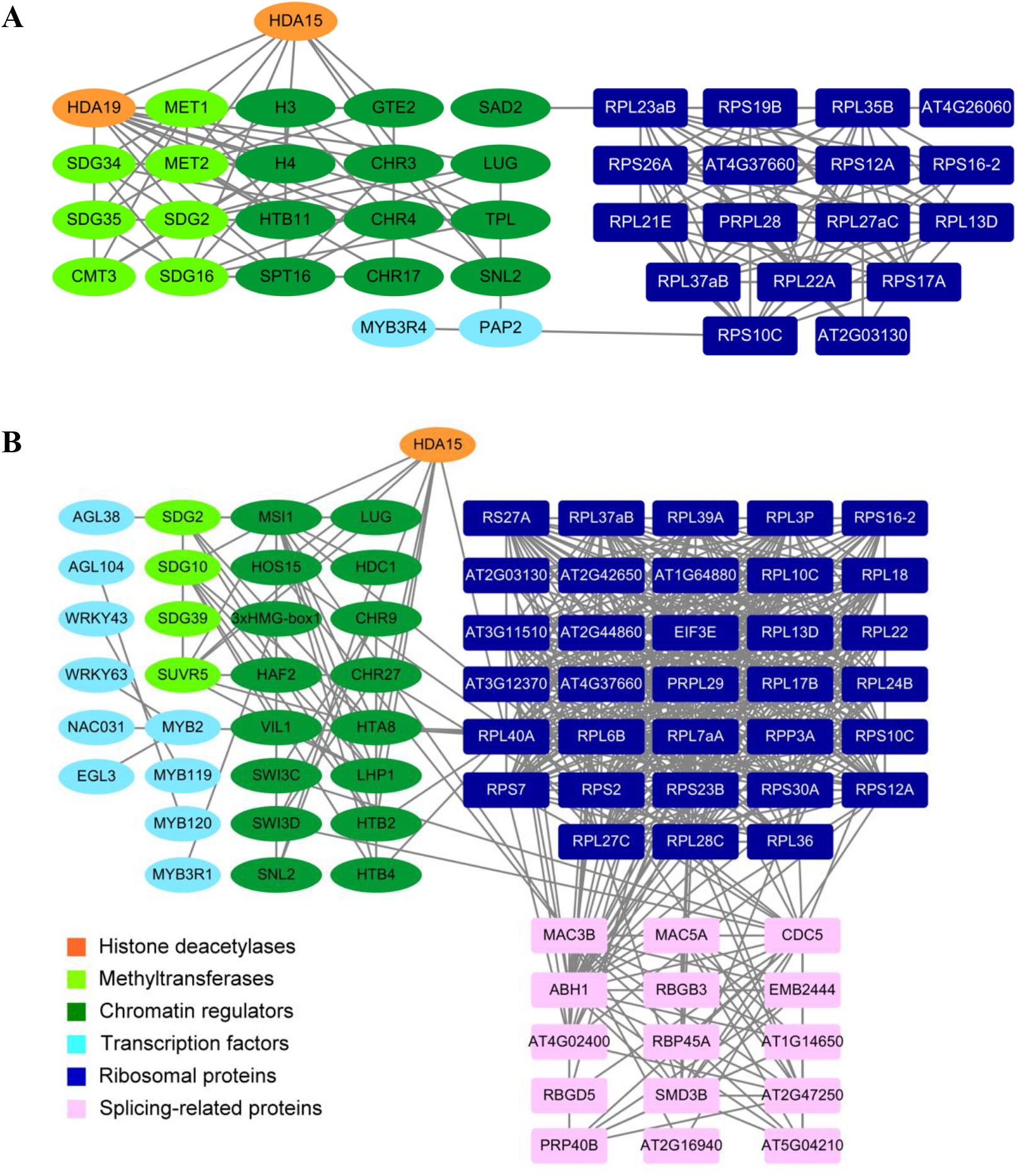
The functional network of HDA15 interacting proteins. The interaction proteins correlation network in mock (A) and ABA treatment. Data were analyzed by STRING and visualized by Cytoscape.

In both mock and ABA treatment, several histone methyltransfreases including SET DOMAIN PROTEIN 16 (SDG16) and SU(VAR)3-9-RELATED PROTEIN 5 (SUVR5) were identified. Under ABA treatment, two WD-40 proteins, MSI1 (MULTICOPY SUPRESSOR OF IRA1(MSI1) and HIGH EXPRESSION OF OSMOTICALLY RESPONSIVE GENES 15 (HOS15), were identified. MSI1 is part of an HDA complex and is involved in ABA signaling (Mehdi et al., 2016). HOS15 is involved in cold stress by affecting the histone acetylation of the stress responsive genes (Zhu et al., 2008). In addition, several transcription factors including MYB2 and WRKY63 were identified under ABA treatment. MYB2 is known to regulate the expression of salt- and dehydration-responsive genes (Abe et al., 2003), and WRKY63 is involved in the regulation of plant responses to ABA and drought stress (Ren et al., 2010). Interestingly, a group of splicing-related proteins including the core subunits of MAC, MAC3B, MAC5A, and CDC5, were identified only in ABA treatment conditions (Fig. 1B).

### HDA15 interacts with MAC3A and MAC3B *in vivo* and ABA enhances the interaction between HDA15 and MAC3B

MAC3A and MAC3B are two core subunits of MAC, which is a conversed complex that is associated with the spliceosome (Palma et al., 2007; Monaghan et al., 2009; Monaghan et al., 2010). Previous studies indicate that MAC3A and MAC3B function redundantly in plant innate immunity (Monaghan et al., 2009) and salt tolerance (Li et al., 2019). Since MAC3B was identified as a HDA15 interacting protein in mass spectrometry analysis, we also analyze whether HDA15 can also interact with MAC3B. The interaction between MAC3A/MAC3B and HDA15 was further confirmed by bimolecular fluorescence complementation (BiFC) assays and coimmunoprecipitation (co-IP) assays. As shown in Fig. 2A, both MAC3A and MAC3B interacted with HDA15 in the nucleus in Arabidopsis protoplasts in BiFC assays.

**Fig. 2.**
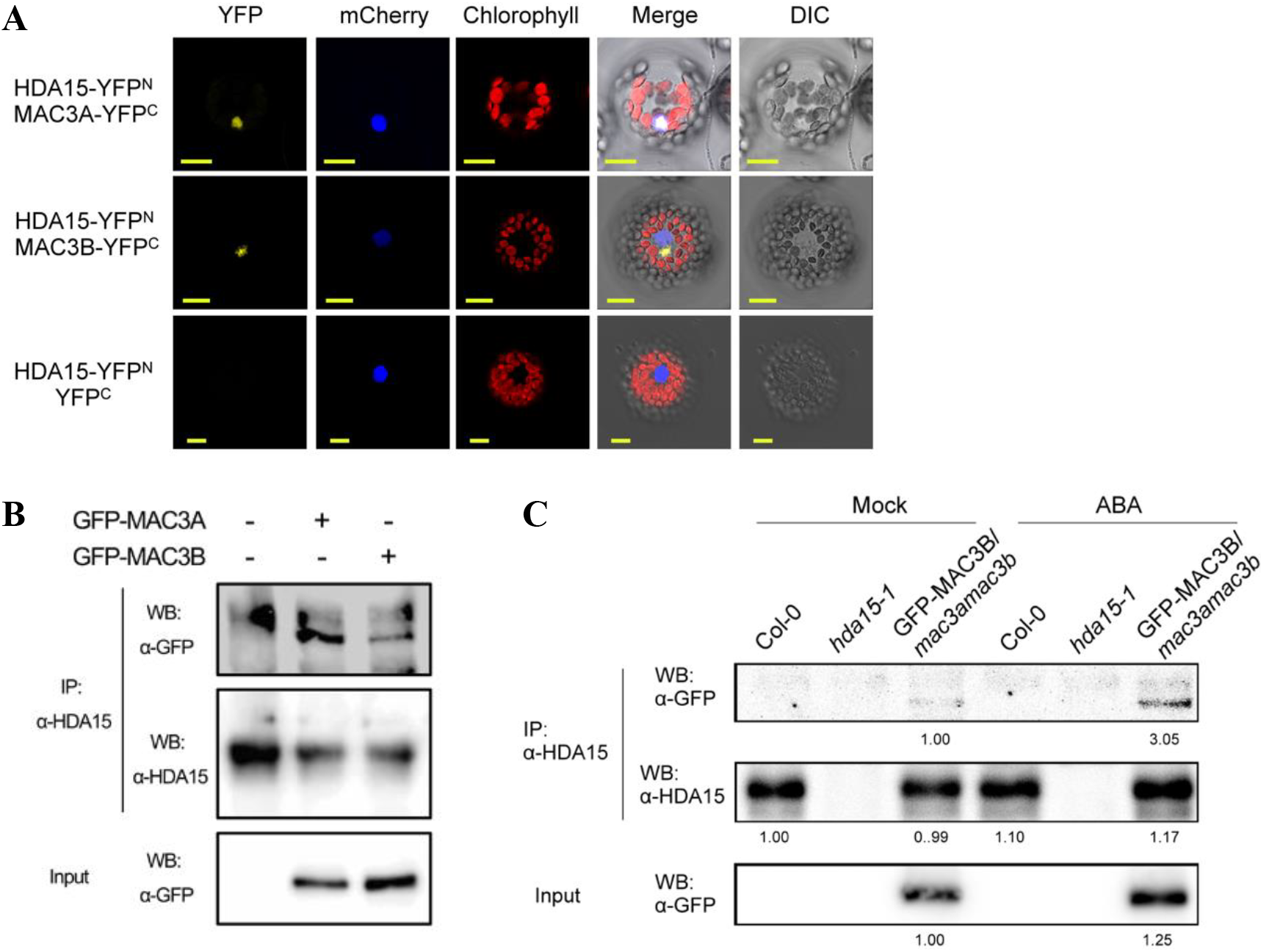
HDA15 interacted with MAC3A and MAC3B. (A) HDA15 fused with the YFP N-terminal fraction (HDA15-YFP^N^) and MAC3A/MAC3B fused with the YFP C-terminal fraction (MAC3A-YFP^C^, MAC3B-YFP^C^) were cotransformed into Arabidopsis protoplasts. BiFC signals were observed by a confocal microscope. Nuclear localization was detected by NLS-mCherry. Scale bar, 10 μm. (B) Co-immunoprecipitation analysis from the transient expression of GFP-MAC3A or GFP-MAC3B in Col-0 Arabidopsis protoplasts. Anti-GFP was used for IP and endogenous HDA15 was detected by anti-HDA15. (C) Co-immunoprecipitation analysis of interaction between HDA15 and MAC3B in *Pro35S:GFP-MAC3B* transgenic plants in mock and ABA treatment. Col-0 and *hda15-1* were used as negative controls of Western blot. Anti-GFP was used for IP, and endogenous HDA15 was detected by anti-HDA15. The numbers below indicate the quantitative results normalized to mock and calculated using ImageJ software.

For co-IP assays, MAC3A or MAC3B fused with GFP (*Pro35S:GFP-MAC3A* or *Pro35S:GFP*-*MAC3B*) were transformed into Arabidopsis protoplasts. An anti-HDA15 antibody was used for immunoprecipitation and an anti-GFP antibody was then used for immunoblot analysis. GFP-MAC3A and GFP-MAC3B proteins could be precipitated by the anti-HDA15 antibody in Arabidopsis protoplasts (Fig. 2B), supporting that MAC3A and MAC3B interact with HDA15 *in vivo*.

Moreover, we also examined the interaction of HDA15 and MAC3B under ABA treatment conditions. The *Pro35S:GFP-MAC3B* transgenic plants expressing GFP-MAC3B driven by the 35S promoter were treated with or without ABA. Although GFP-MAC3B was coimmunoprecipitated by endogenous HDA15 in both mock and ABA-treated conditions, the interaction between HDA15 and MAC3B was greatly enhanced by ABA treatment (Fig. 2C), indicating that ABA promote the interaction between HDA15 and MAC3B.

### *mac3a/3b* plants are hyposensitive to abiotic stress and ABA

Since HDA15 is involved in abiotic stress and ABA responses (Lee and Seo, 2019), we also investigated whether MAC3A and MAC3B are also involved in abiotic stress and ABA responses by comparing the phenotypes of the single mutants *hda15*, *mac3a* and *mac3b* as well as the double mutants *mac3a/hda15*, *mac3b/hda15* and *mac3a/mac3b*. After salt stress treatment, about 50-60% *hda15-1* and *mac3a/mac3b* plants survived, compared to the survival rate of 30% Col-0 wild type (WT) plants (Fig. 3A, B). The water loss rates of detached leaves in *hda15-1* and *mac3a/3b* plants were also higher compared to WT (Fig. 3C). Furthermore, *hda15* and *mac3a/3b* had higher germination rates on the media containing ABA (Fig. 3D). These results indicate that MAC3A and MAC3B are also involve in abiotic stress and ABA response similar to HDA15. No obvious phenotype difference was observed in the *mac3a* and *mac3b* single mutants compared to WT. Furthermore, the phenotypes of *mac3a/hda15* and *mac3b/hda15* double mutants were similar to that of the *hda15* single mutant in salt tolerance, transpiration rates, and seed germination (Fig. 3). These observations support the notion that MAC3A and MAC3B function redundantly in abiotic stress and ABA responses.

**Fig. 3.**
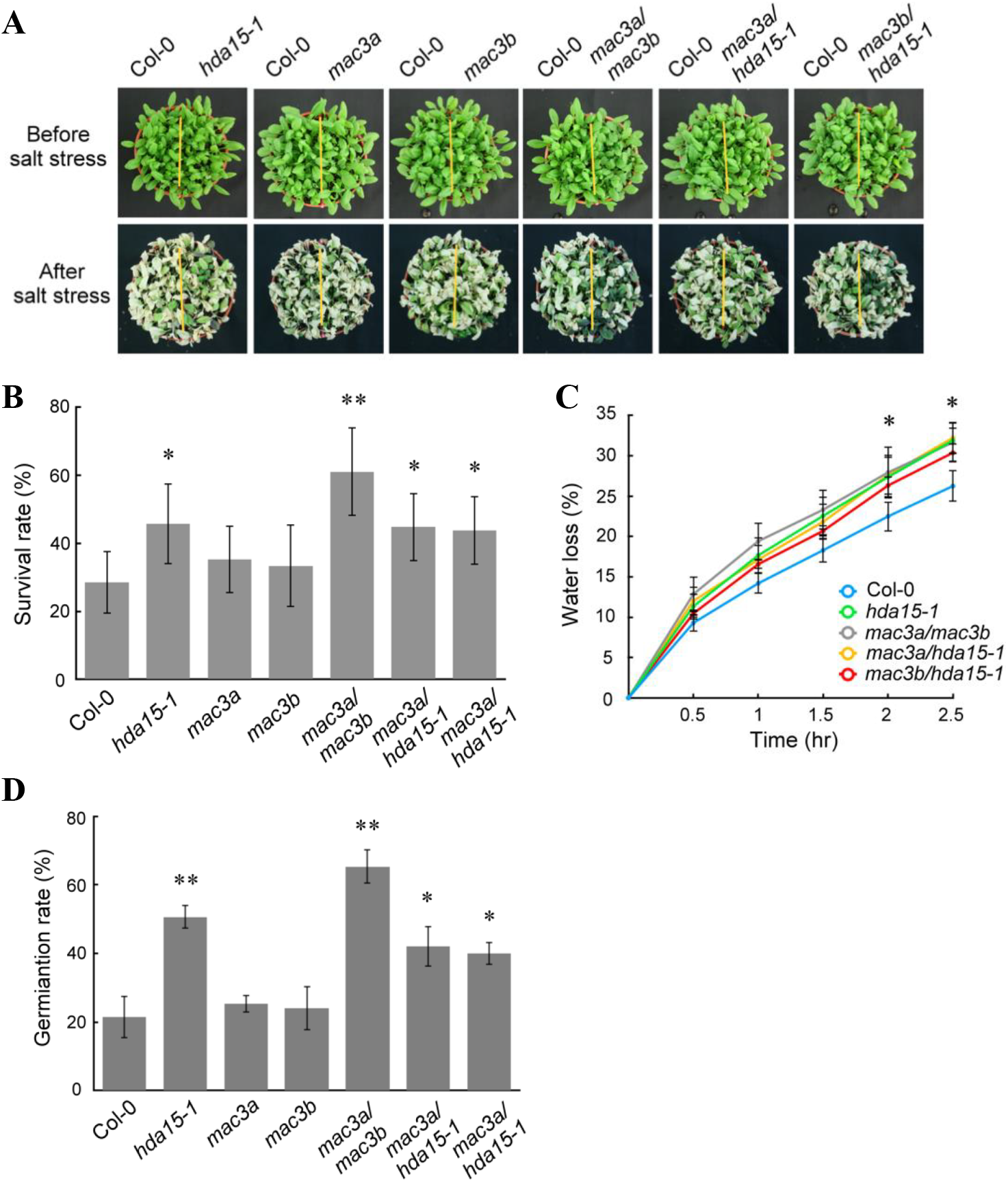
The phenotypes of *mac3a*, *mac3b*, *mac3a/3b*, *mac3a/hda15*, and *mac3b/hda15* in response to salt stress and ABA. (A) The phenotype of three-week-old plants irrigated with 300mM NaCl solution for two weeks. (B) Survival rates of three-week-old plants irrigated with 300mM NaCl solution for two weeks. Error bars represent SD of triplicates (*t* test, *p<0.05, **p<0.01). (C) Water loss from detached leaves in Col-0, *hda15-1*, *mac3a*, *mac3b*, *mac3a/mac3b*, *mac3a/hda15*, and *mac3b/hda15* plants. Values are the mean of the percentage of leaf water loss ± SD (*t* test, *p<0.05, *n*= 20). (D) Germination rates of Col-0, *hda15-1*, *mac3a*, *mac3b*, *mac3a/mac3b*, *mac3a/hda15*, and *mac3b/hda15* seeds germinated on the media containing 2 μM ABA for 3 days. Experiments were performed in triplicate and repeated at three times with similar results. Values are shown as means ± SD (*t* test, *p<0.05, **p<0.01; *n*=50).

### Transcriptome analysis of *hda15-1* and *mac3a/mac3b* mutants

To further characterize the role of HDA15, MAC3A, and MAC3B in ABA responses, 10-day-old Col-0, *hda15-1*, and *mac3a/3b* plants treated with or without (mock) ABA were used for RNA-seq analysis. Compared with WT in mock conditions, 78 and 705 genes displayed higher transcript levels (fold change ≥ 1.5 with *P* < 0.05) in *hda15-1* and *mac3a/mac3b* plants, respectively (Supplemental Data Set 2, 3). Among these upregulated genes from *hda15-1* and *mac3a/mac3b*, 9 of them (Fisher’s exact test*, P* <8.9e-06) were upregulated in both *hda15-1* and *mac3a/mac3b* (Fig. 4A, Supplemental Data Set 4). We also identified 115 and 965 downregulated genes in *hda15-1* and *mac3a/mac3b* plants, respectively (Supplemental Data Set 2, 3). Among these downregulated genes, 18 of them (Fisher’s exact test*, P* <1.8e-10) overlapped (Fig. 4A, Supplemental Data Set 4). The expression of many ABA-responsive genes including NADPH-oxidase *RbohI*, *ABI5 BINDING PROTEIN 4* (*AFP4*), and *BREVIS RADIX* (*BRX*) was impaired in *hda15* plants. In addition, the expression of *PYL7* encoding an ABA receptor and some ABA-responsive genes including *RD22*, *COLD-REGULATED 47* (*COR47*), and *POTASSIUM CHANNEL IN ARABIDOPSIS THALIANA 1* (*KAT1*) was affected in *mac3a/mac3b* plants. Furthermore, the salt stress-responsive genes *GLUTATHIONE S-TRANSFERASE 7* (*GSTF7*) and *LATE EMBRYOGENESIS ABUNDANT 7* (*LEA7*) were affected in both *hda15* and *mac3a/mac3b* plants in the mock condition (Supplemental Data Set 4).

**Fig. 4.**
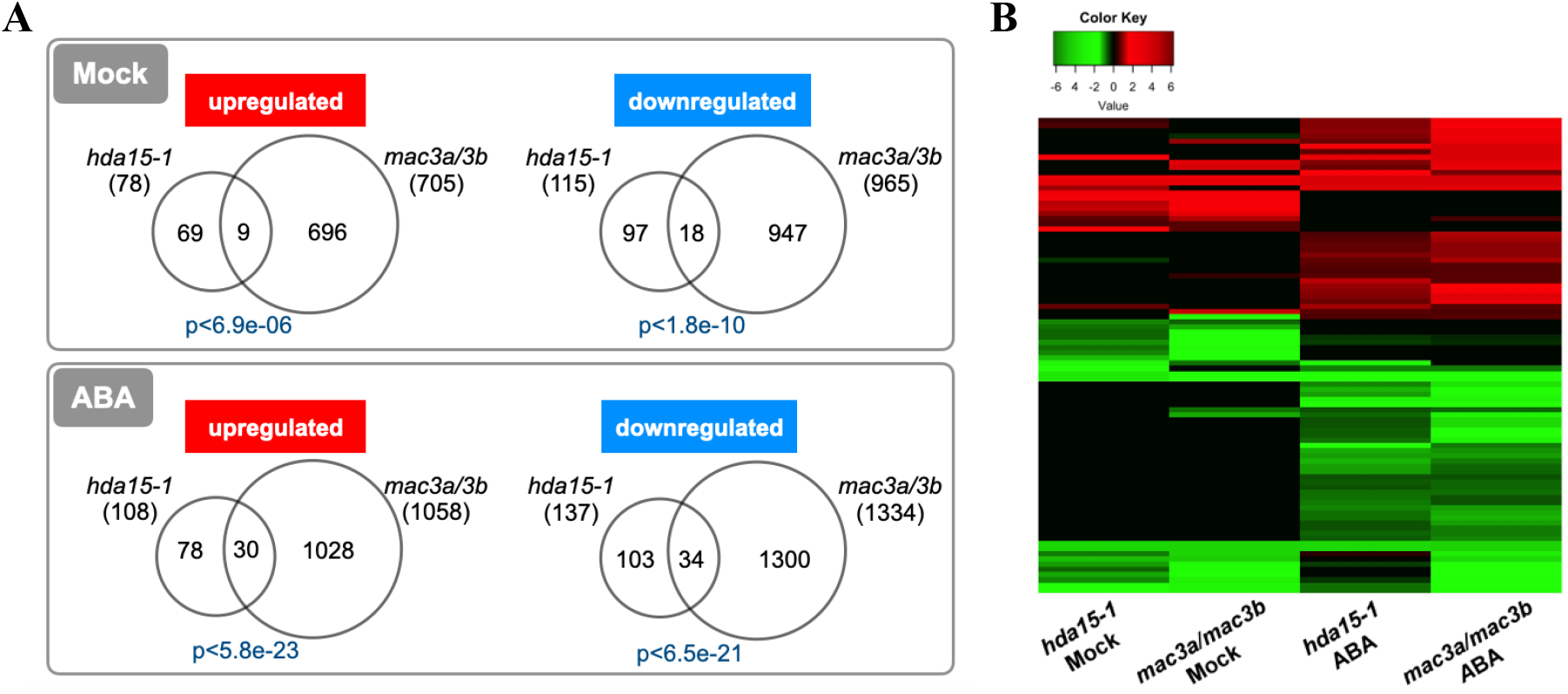
Genome-wide expression analysis in Col-0, *hda15-1*, and *mac3a/3b* by RNA-seq. (A) Venn diagram of the number of genes that displayed differential expression in *hda15-1* and *mac3a/mac3b* compared with Col-0 in mock or ABA treatment. RNA-seq analysis was performed using 10-d-old Col-0, *hda15-1*, and *mac3a/3b* in mock or treated with 50 μM ABA for 3 hr. The p-value of overlapped genes was determined by the Fisher’s exact test. (B) The heatmap shows the fold changes of the coregulated genes in *hda15-1* and *mac3a/mac3b*. The fold changes were log_2_ transformed.

Under ABA treatment, 108 and 1058 genes were upregulated (fold change ≥ 1.5 with *P* < 0.05) in *hda15-1* and *mac3a/mac3b*, respectively (Supplemental Data Set 2, 3), with 30 genes (Fisher’s exact test*, P* <5.8e-23) overlapped (Fig. 4A, Supplemental Data Set 4). 137 and 1334 genes were downregulated genes in *hda15-1* and *mac3a/mac3b*, respectively (Supplemental Data Set 2, 3), with 34 genes (Fisher’s exact test*, P* <6.5e-21) overlapped (Fig. 4A, Supplemental Data Set 4). The heatmap analysis showed that the overlapped genes in *hda15-1* and *mac3a/mac3b* displayed similar patterns (Fig. 4B). The transcriptome analysis showed that the expression of *ATP-BINDING CASSETTE G40* (*ABCG40*) encoding an ABA transporter was downregulated in *hda15* plants. The expression of the ABA signaling pathway genes such as *SNRK2.2*, *ABA INSENSITIVE 5* (*ABI5*) and *ABSCISIC ACID RESPONSIVE ELEMENT-BINDING FACTOR 1* (*ABF1)*, and a key enzyme gene in ABA biosynthesis, *NINE-CIS-EPOXYCAROTENOID DIOXYGENASE 3* (*NCED3*), was affected in *mac3a/mac3b* plants. Furthermore, the salt stress-responsive genes, *NA^+^/H^+^* (*SODIUM HYDROGEN*) *EXCHANGER 3* (*NHX3*) and *HEAT SHOCK PROTEIN 22.0* (*HSP22.0*), and the cold stress-responsive gene *EPITHIOSPECIFIER MODIFIER 1* (*ESM1*) were affected in both *hda15* and *mac3a/mac3b* plants under ABA treatment (Supplemental Data Set 4).

Together, our genome-wide expression analyses suggest that HDA15 and MAC3A/MAC3B coregulate a subset of genes involved in ABA responses. In addition, a large number of differentially expressed genes that are not overlapped in *hda15-1* and *mac3a/mac3* plants indicate that HDA15 and MAC3A/MAC3B also function independently in gene regulation.

### HDA15 and MAC3A/MAC3B are involved in alternative splicing in ABA responses

MAC3A and MAC3B are the core subunits of MAC, which plays a critical role in pre-mRNA splicing (Palma et al., 2007; Monaghan et al., 2009; Monaghan et al., 2010). We further explored whether HDA15, MAC3A, and MAC3B coregulate pre-mRNA splicing to mediate ABA responses by analyzing genome-wide alternative splicing. AS events can be classified into five types: alternative donor (AltD), alternative acceptor (AltA), alternative donor and acceptor (AltD/A), exon skipping (ES), and IR. Compared with Col-0 in mock treatment, 3,240 and 20,582 novel IR events were identified in *hda15-1* and *mac3a/3b*, respectively (Fig. 5A, Supplemental Data Set 5). In ABA treatment, IR events were dramatic increased to 5,462 and 31,311 in *hda15-1* and *mac3a/3b*, respectively (Fig. 5A). Over 90% of AS events were IR (Supplemental Data Set 5~7). Since IR is the major mode of AS events, we further analyzed the patterns of the IR events by monitoring relative IR levels. The IR level was defined as the read coverage depth of an intron divided by that of the two neighboring exons. *hda15-1* and *mac3a/3b* had higher IR levels than Col-0 both in mock and ABA treatment (Fig. 5B).

**Fig. 5.**
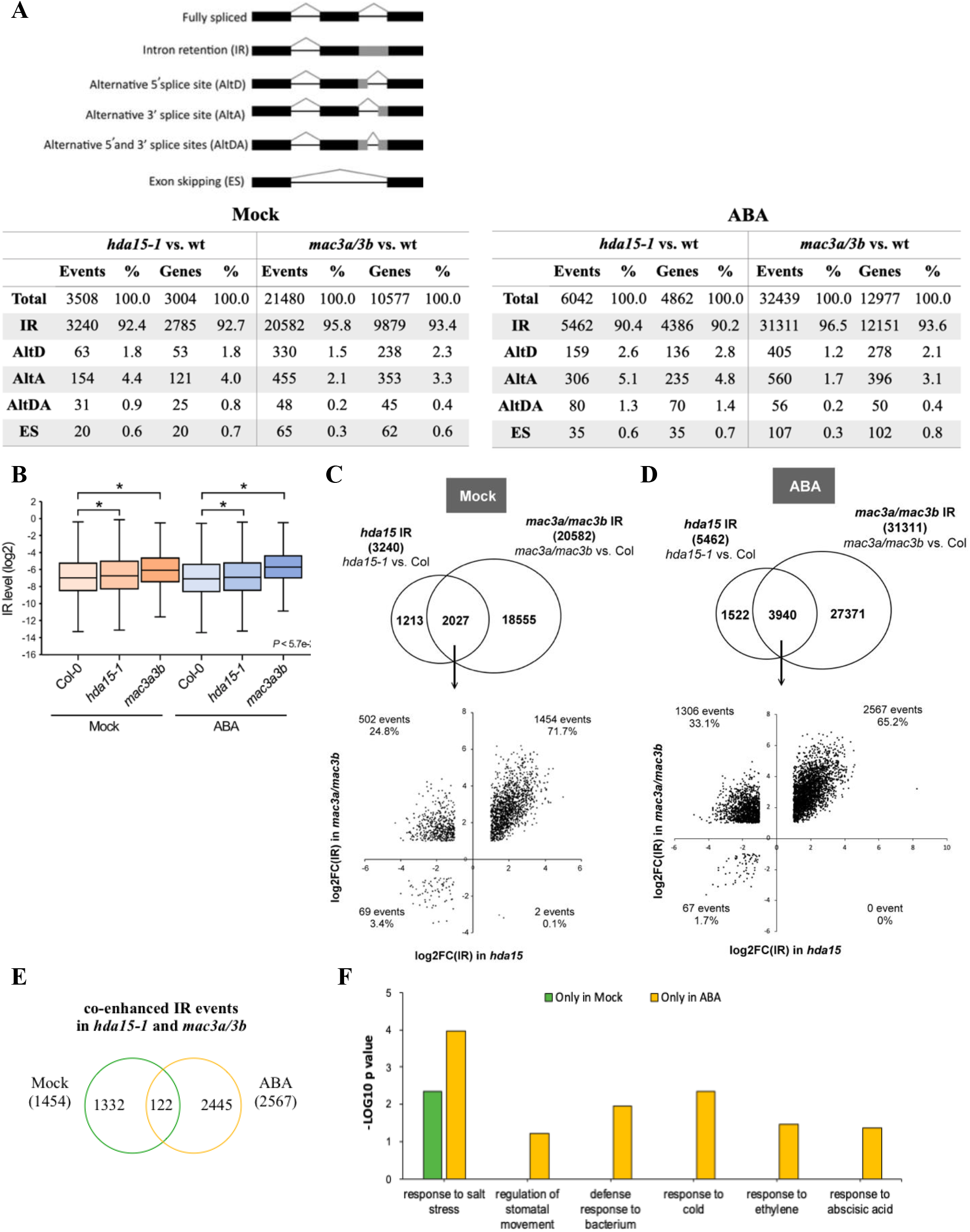
Genome-wide splicing analysis and intron retention levels of *hda15-1* and *mac3a/mac3b* compared with Col-0 in mock and ABA treatment. (A) Alternative splicing events in *hda15-1*, and *mac3a/mac3b* compared with Col-0 in mock and ABA treatment. (B) Box plot of total IR levels. The log2 IR levels of alternative splicing events (p<0.05 and fold change >2, *t* test) were plotted. (C) HDA15, MAC3A, and MAC3B coregulated intron retention events. Venn diagram comparing IR events regulated by HDA15, MAC3A, and MAC3B (up panel). Scatter Chart showed the IR trend in *hda15* and *mac3a/3b* in mock (down panel). The numbers mean IR events and the percentage in the quadrant. (D) Venn diagram comparing IR events regulated by HDA15, MAC3A, and MAC3B (up panel). Scatter Chart showed the IR trend in *hda15-1* and *mac3a/3b* in ABA treatment (down panel). The numbers mean IR events and the percentage in the quadrant. (E) Venn diagram shows co-enhanced IR events regulated by HDA15, MAC3A, and MAC3B in mock and ABA. 1332 and 2567 co-enhanced IR events were specific in mock and ABA treatment, respectively. (F) GO analysis of co-enhanced IR in *hda15-1* and *mac3a/3b*. The specific IR events in ABA treatment were highly enriched with ABA and stress related GO terms.

We further analyzed the coregulated IR events in *hda15-1* and *mac3a/3b*. The Venn diagram showed that among the 2,027 coregulated IR events in mock treatment, 1,454 of them (71.7 %) were co-enhanced as shown in the quadrant plot (Fig. 5C). In ABA treatment, among the 3,940 coregulated events, 2,567 of them (65.2 %) showed a similar pattern of enhances in *hda15-1* and *mac3a/3b* (Fig. 5D). In the co-enhanced IR events, 1,454 and 2,567 events were specifically occurred in mock and in ABA treatment, respectively (Fig. 5E). The Gene Ontology analysis showed that “response to salt stress” was enriched in both mock and ABA treatment (Fig. 5F). Interestingly, ABA- and stress-related GO were highly increased in ABA treatment (Fig. 5F). Taken together, these results indicate that HDA15 and MAC3A/MAC3B coregulate intron retention of ABA- and stress-related genes in ABA response.

Compared to mock treatment, 5,763 and 5,497 AS events were identified after ABA treatment in Col-0 and *hda15-1*, respectively. There were 5,271 IR events in Col-0, which account for 91.5% of all AS events. There were 5,008 IR events in *hda15-1*, which also account for 91.1% of all AS events. In particular, a large number of AS (15,896) and IR (15,385) events were identified in *mac3a/3b* after ABA treatment (Fig. 6A, Supplemental Data Set 8). We further analyzed whether loss of function of HDA15, MAC3A, or MAC3B causes IR changes in ABA responses by analyzing relative IR levels. After ABA treatment, the IR levels of Col-0 and *hda15-1* were slightly decreased compare to mock treatment. However, the IR level of *mac3a/mac3b* was increased after ABA treatment (Fig. 6B), indicating that MAC3A and MAC3B are required for the genome-wide alternative splicing in response to ABA.

**Fig. 6.**
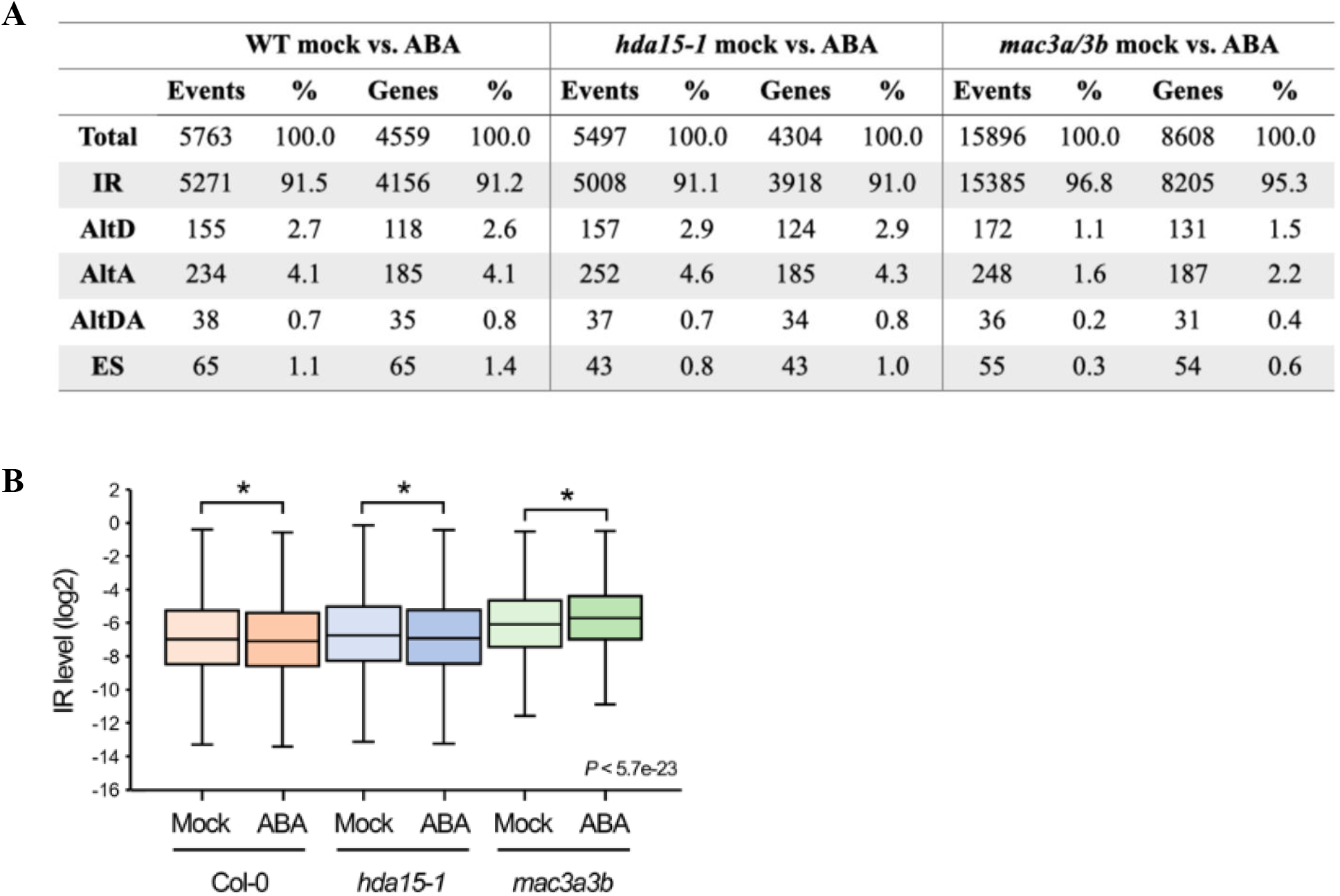
Genome-wide pre-mRNA splicing analysis in ABA treatment by RNA-seq. (A) Novel alternative splicing events in Col-0, *hda15-1*, and *mac3a/mac3b* with ABA treatment. (B) Box plot of total IR level. The log2 IR levels of alternative splicing events (p<0.05 and fold change >2, *t*-test) were plotted.

### HDA15 and MAC3A/MAC3B affect ABA-responsive IR

To further uncover the involvement of HDA15 and MAC3A/MAC3B in ABA-responsive alternative splicing, we defined those IR events with more than two-fold enrichment after ABA-treatment in Col-0 as ABA-responsive IR events, and 5,271 ABA-responsive IR events were identified (Fig. 6A, Supplemental Data Set 8). We compared the IR levels of the ABA-responsive IR events in Col-0 with those in *hda15-1* and *mac3a/3b.* The IR level was slightly increased in Col-0, *hda15-1,* and *mac3a/3b* when treated with ABA (Fig. 7A). We further classified the ABA-responsive IR events into the enhanced or reduced events. In the number of enhanced ABA-responsive IR events, Col-0 had a 2.9-fold increase after ABA treatment, whereas *hda15-1* and *mac3a/mac3b* only had 1.5- and 1.4-fold increases, respectively (Fig. 7B). In the number of reduced ABA-responsive IR events, Col-0 had a 3.2-fold decrease, whereas *hda15-1* and *mac3a/mac3b* had almost no changes. (Fig. 7C). These data indicate that HDA15 and MAC3A/MAC3B play important roles in modulating ABA-responsive IR, especially the reduced ABA-responsive IR events.

**Fig. 7.**
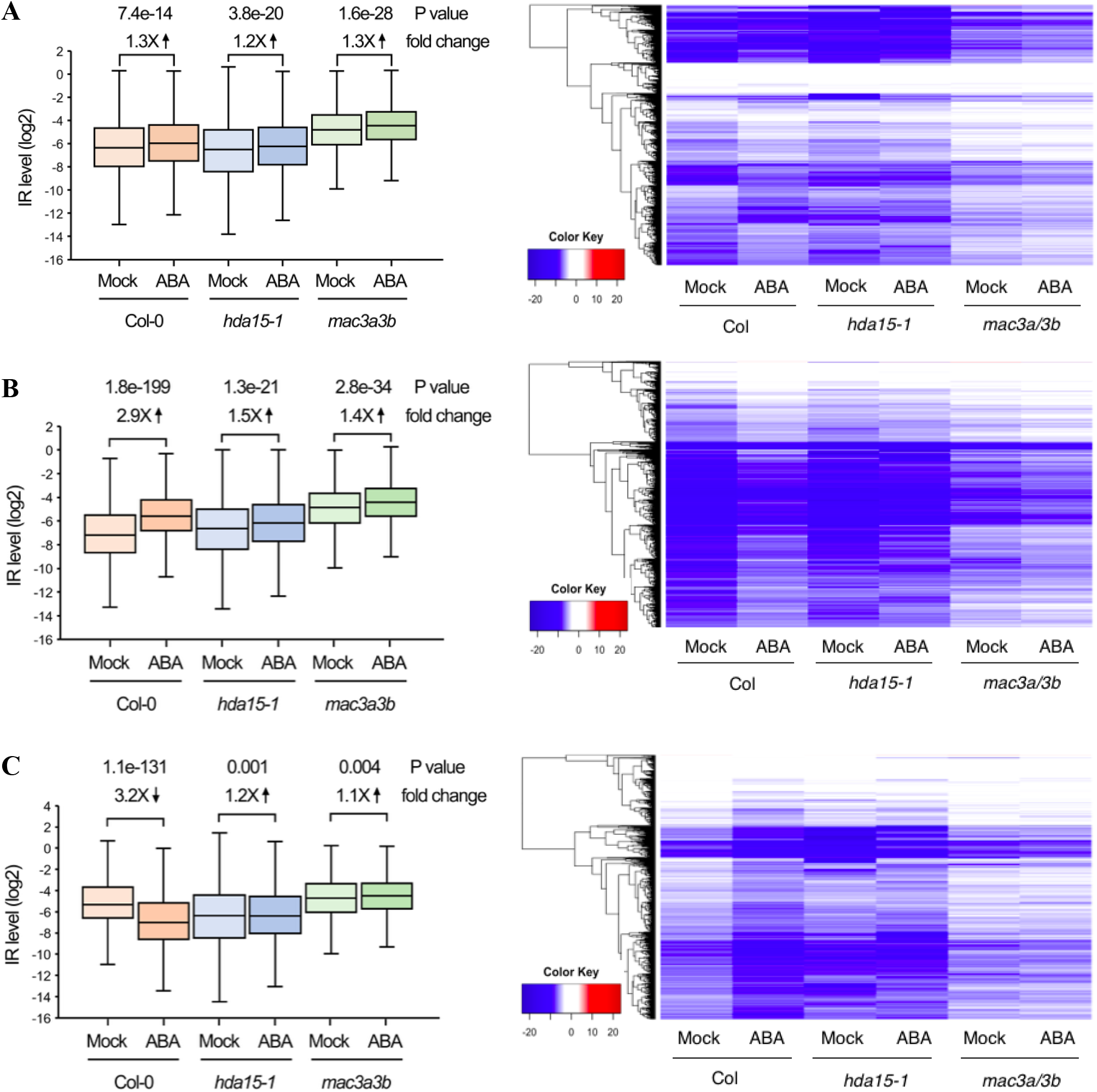
ABA-responsive IR levels in Col-0, *hda15-1*, and *mac3a/3b* in mock and ABA treatment. (A) ABA-responsive IR levels (left panel) and heatmap (right panel). (B) Box plot of enhanced ABA-responsive IR levels (left panel) and heatmap (right panel). (C) Box plot of reduced ABA-responsive IR levels (left panel) and heatmap (right panel). The log2 IR levels of alternative splicing events (p<0.05 and fold change >2, *t* test) were plotted.

The effect of ABA on IR of three selected ABA- and stress-related genes, *SNRK2.6*, *NTAQ1* (*GLN-SPECIFIC AMINO-TERMINAL* (*NT*)*-AMIDASE*), and *EDM1* (*ENHANCED DOWNY MILDEW 1*), in *hda15* and *mac3a/mac3b* was examined by qRT-PCR. The IR form of *SNRK2.6* was increased in *hda15* and *mac3a/mac3b* compared to Col-0, and ABA further increased its level (Fig. 8A, D). The IR forms of *NTAQ1* and *EDM1* were increased in *mac3a/mac3b* but not in *hda15* in mock treatment. However, the levels of these IR forms were induced in both *had15* and *mac3a/3b* in ABA treatment, which is consistent with the RNA-seq data (Fig. 8B, C, E, F). These results indicate that HDA15 and MAC3A/MAC3B coregulate intron retention of ABA-related genes, and ABA enhances the IR events regulated by HDA15 and MAC3A/MAC3B.

**Fig. 8.**
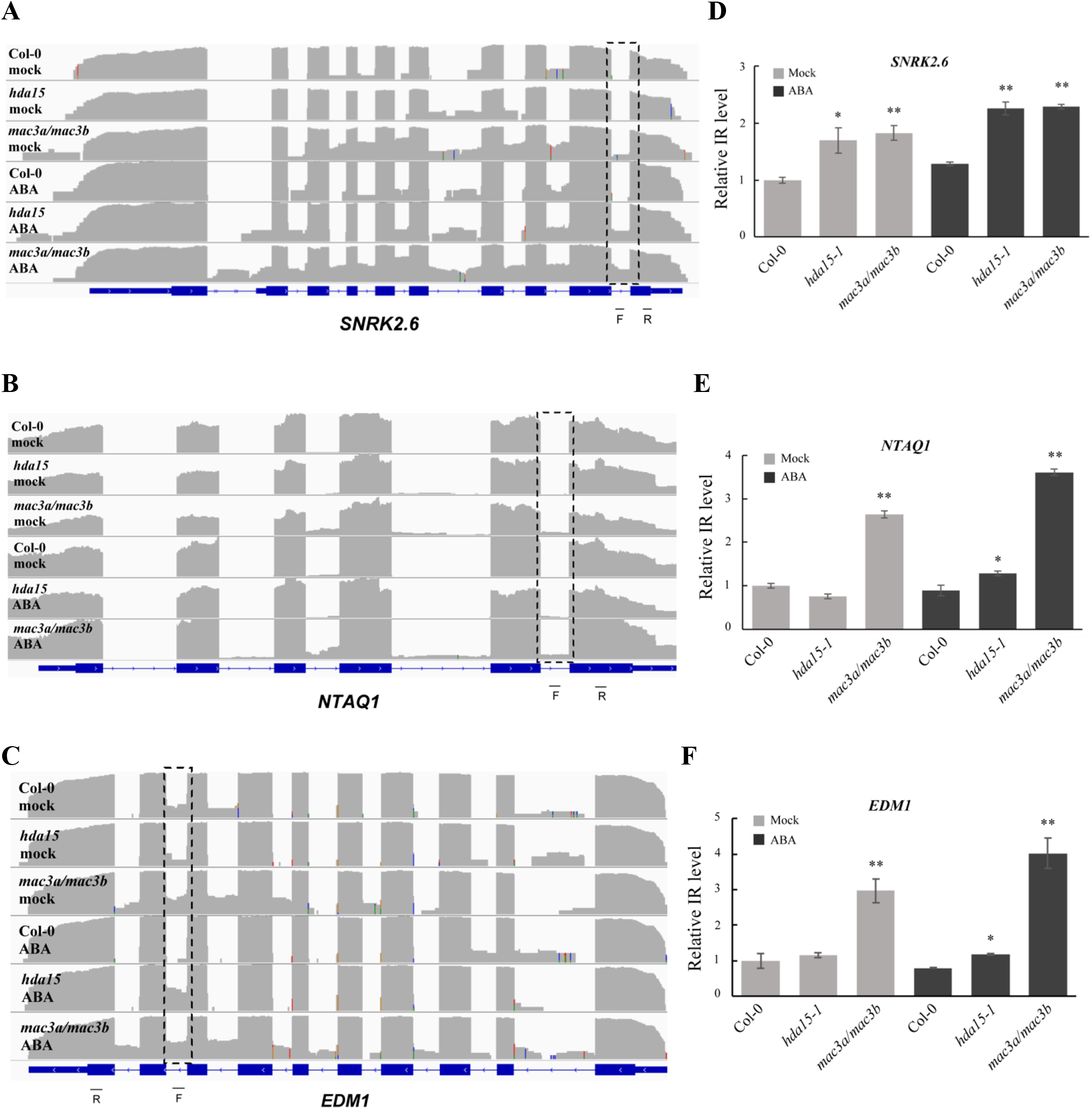
ABA-responsive IR events defects in *hda15-1* and *mac3a/mac3b* in ABA treatment. (A) to (C) Read mapping of ABA-responsive IR events visualized in the Integrative Genomics Viewer (https://software.broadinstitute.org/software/igv/). Dashed boxes indicate the regions of IR events. F and R indicate the forward and reversed primer using in quantitative RT-PCR. (D) to (F) The levels of ABA-responsive IR events in Col-0, *hda15-1*, and *mac3a/mac3b* plants in mock and ABA treatment analyzed by quantitative RT-PCR. The relative expression was normalized with Col-0. *UBQ* was used as an internal control. Error bars represent SD.

HDAs remove acetyl groups from ε-N-acetyl lysine residues of histones. We further investigated whether HDA15 and MAC3A/MAC3B affect the histone acetylation levels near the retained intron by chromatin immunoprecipitation (ChIP) assays. We examined the levels of H3K9ac approximately 250 bp upstream of the retained intron (P1), approximately 250 bp downstream of the retained intron (P2), and about 800 bp downstream from P2 as a control (P3). The H3K9ac levels near the 9^th^ intron of *SNRK2.6*, the 5^th^ intron of *NTAQ1*, and the 8^th^ intron of *EDM1* (P1 and P2) were increased in *hda15* and *mac3a/mac3b* when treated with ABA compared with Col-0. In contrast, there was no difference in the H3K9ac levels of the P3 regions, which are located far away from the retained introns (Fig. 9). These results indicate that HDA15 and MAC3A/MAC3B affect the histone acetylation level of the regions near ABA-responsive IRs in response to ABA.

**Fig. 9.**
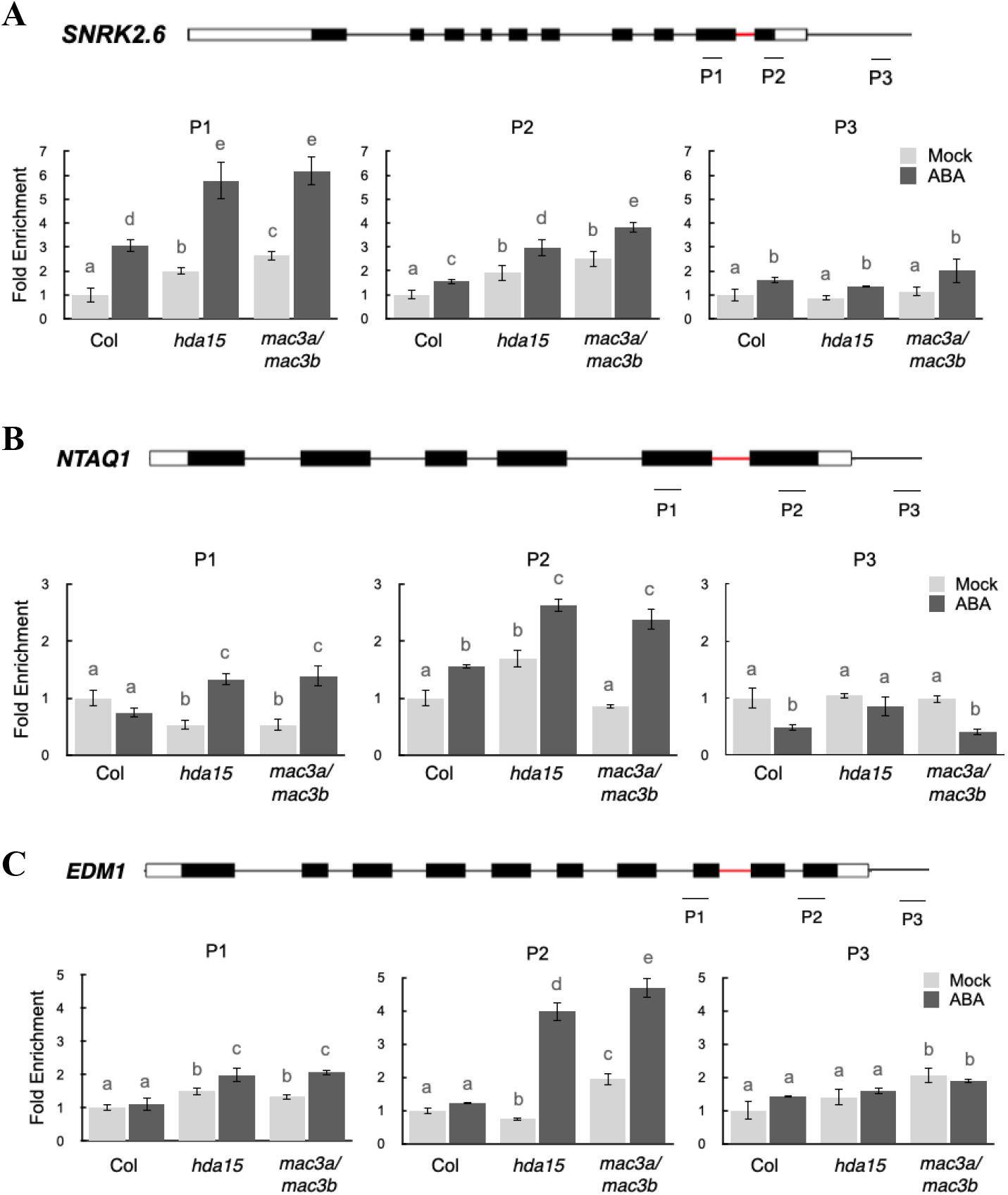
H3K9ac levels of the retained introns of *SNRK2.6*, *NTAQ1*, and *EDM1* in *hda15* and *mac3a/mac3b*. Schematic diagrams of *SNRK2.6*(A), *NTAQ1*(B), and *EDM1*(C) genes with amplicons (indicated as P1 to P3) used for ChIP-qPCR were shown. Open boxes represent UTRs, and black boxes represent exons. The red bars indicate the introns that were retained in *hda15-1* and *mac3a/3b* in RNA-seq. 10-day-old Col-0, *hda15-1*, and *mac3a/mac3b* treated with or without (mock) ABA were used for ChIP-qPCR using an anti-H3K9ac antibody. The amounts of DNA relative to Col-0 in mock were normalized to *ACTIN7*. Different letters above the error bars indicate statistically significant difference (*P* < 0.05, two-way ANOVA with Tukey HSD Test).

## Discussion

It has been reported that MYB96 recruits HDA15 to suppress *RHO GTPASE OF PLANTS* (*ROP*) genes by histone deacetylating in the presence of ABA(Lee and Seo, 2019). To further investigate the function of HDA15 in ABA responses, we used immunopurification coupled with mass spectrometry-based proteomics to identify the HDA15 interacting proteins. We found that HDA15 can interact with many transcription factors, chromatin remodeling proteins, and ribosomal proteins. HDA15 copurified with the WD40-repeat proteins MSI1 and HOS15. MSI1 is part of an HDA complex and loss of function of MSI1 causes upregulation of ABA receptor genes and hypersensitivity of ABA-responsive genes (Mehdi et al., 2016). The WD40-repeat protein HOS15 shares sequence similarity with human TBL1 (transducin-beta like protein 1), which is a component of the repressor complex that associates with HDAs (Zhu et al., 2008). The reduction of HOS15 results in a cold hypersensitive phenotype in Arabidopsis (Zhu et al., 2008). HDA15 may form an HDA complex with MSI1 and HOS15 to mediate ABA responses in Arabidopsis.

Recent studies have revealed that alternative splicing regulates stress responses through targeting the ABA signaling pathway. RNA-BINDING PROTEIN 25 (RBM25) binds to the *PP2C HYPERSENSITIVE TO ABA1* (*HAB1*) pre-mRNA, resulting in two splicing variants, *HAB1.1* and *HAB1.2*. HAB1.1 interacts with SnRK2.6/OST1 and inhibits its kinase activity to turn off the ABA signaling (Wang et al., 2015). SNW/SKI-INTERACTING PROTEIN (SKIP), an essential spliceosomal component and transcriptional coregulator, interacts with MAC3A and MAC3B to mediate the pre-mRNA splicing of abiotic stress-related genes, thereby affecting environmental fitness (Li et al., 2019). We found that HDA15 also interacts with the core subunits of the MAC complex, MAC3A and MAC3B, and modulates ABA-responsive IR events, suggesting that HDA15 and MAC3A/MAC3B may co-regulate ABA responses and stress adaption through alternative splicing.

The involvement of histone modifications in splicing regulation has been reported in yeasts and animals. In mammals, H3K36me3 on the alternatively splicing regions is recognized by MORF-Related Gene 15 (MRG15), which recruits the PTB splicing factor to promote exon skipping. In contrast, the low level of H3K36me3 causes exon inclusion (Luco et al., 2010). In Arabidopsis, the H3K36me3 methyltransferase mutants, *sdg8* and *sdg26*, disturb the alternative splicing of the flowering regulator gene *FLM* and the circadian clock gene *PRR7* (Pajoro et al., 2017). The MRG15 homologs MRG1 and MRG2 in Arabidopsis also bind to H3K36me3 (Xu et al., 2014). Moreover, the PTB homologs in Arabidopsis, PTB1 and PTB2, mediate exon inclusion within *FLM* (Ruhl et al., 2012). In addition to H3K36me3, H3K9 and H3K4 methylations also affect alternative splicing. H3K9me3 is recognized by the heterochromatin 1 protein HP1*r*, leading to a reduce transcriptional elongation rate and exon inclusion in human (Saint-André et al., 2011). In *Drosophila*, H3K9me3 and HP1 interact to recruit heterogeneous nuclear ribonucleoparticles (hnRNPs), which repress splicing by directly antagonizing the recognition of splice sites (Piacentini et al., 2009). In human, H3K4me3 is recognized by the chromatin-adaptor protein CHD1, recruiting spliceosomes to human cyclin D1 pre-mRNA (Sims et al., 2007). The association of HDAs with histone methyltransferases has been reported in plants. Arabidopsis HDA6 interacts with the histone methyltransferases SUVH4/5/6 and coregulates transposons through histone H3K9 methylation and H3 deacetylation (Yu et al., 2017). In the present study, we found that HDA15 also interacts with several histone methyltransferases including SDG16 and SUVR5. It has been reported that SDG16 can bind to ABA-HYPERSENSITIVE GERMINATION 3 (AHG3) and trimethylated H3K4 (Liu et al., 2018), whereas SUVR5 mediates H3K9me2 deposition (Caro et al., 2012). Further research is needed to investigate the function of HDA15 interacting with histone methyltransferases in the regulation of alternative splicing.

In yeast, spliceosome assembly is strongly coupled with histone modifications including acetylation and deacetylation. The yeast histone acetyltransferase Gcn5 is required for the association of U2 snRNP to the splicing branchpoint (Gunderson and Johnson, 2009). Mutation or deletion of Gcn5-targeted histone H3 residues leads to the intron accumulation of the Gcn5 target genes (Gunderson et al., 2011). In yeast, Prp19 is a core component of the Prp19 complex (Prp19C), also known as NineTeen complex (NTC), which functions in splicing and stabilizes U5/U6 snRNP in the spliceosomal complex (Chan et al., 2003). The deletion of the yeast HDACs Hos3 and Hos2 reduces the recruitment of Prp19 to pre-mRNA (Gunderson et al., 2011). MAC3A and MAC3B are the plant homologs of Prp19 (Monaghan et al., 2009). The interaction of MAC3A and MAC3B with HDA15 indicates they are functionally associated. We found that the H3K9 acetylation levels of *SNRK2.6*, *EDM1*, and *NTAQ1* are increased in both in *hda15* and *mac3a/mac3b* mutants, indicating that MAC3A/MAC3B may affect the histone acetylation level of their target genes by interacting with HDA15.

In conclusion, we propose a model of how HDA15 and MAC3A/MAC3B interaction is involved in ABA response through alternative splicing. HDA15 deacetylates the ABA-responsive genes and acts coordinately with MAC3A/MAC3B to regulate alternative splicing in ABA responses (Fig. 10).

**Fig. 10.**
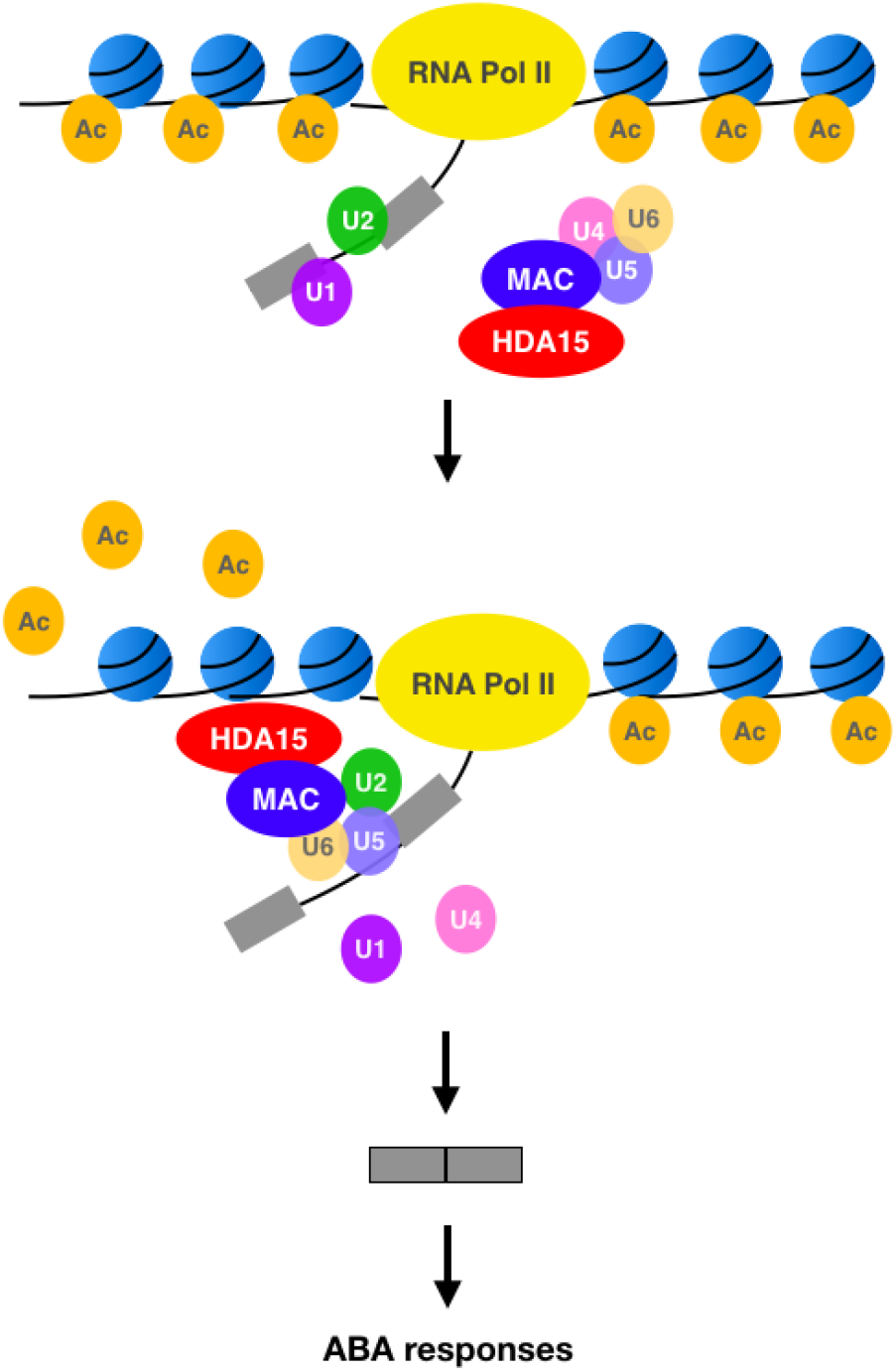
HDA15 interacts with MAC3A and MAC3B and is involved in alternative splicing. HDA15 interacts with MAC3A and MAC3B, two subunits of the MAC complex, which associates with the assembling spliceosome with U4/U5/U6 tri-snRNP. The HDA15-MAC complex is recruited to the splicing site and releases U1 and U4. At the same time, the acetyl groups are removed by HDA15 and introns are spliced out to process mature mRNA.

## Methods

### Plant materials and growth conditions

The T-DNA insertion mutants, *hda15-1* (SALK_004027), *mac3a* (SALK_089300) and *mac3b* (SALK_050811), were described previously (Monaghan *et al.*, 2009; Liu *et al.*, 2013) and all of them are in the Col-0 background. Arabidopsis (*Arabidopsis thaliana*) plants were grown under long-day conditions (16 hr light/ 8 hr dark cycle) at 23°C after seeds were subjected to a 3-day stratification period. To generate *Pro35S:MAC3B-GFP* transgenic lines, MAC3B complementary DNA (cDNA) was cloned into the pK7WGF2 binary vector. The transgenic plants were generated using the floral dip method (Clough & Bent, 1998).

### Liquid chromatography-tandem mass spectrometry (LC-MS/MS)

10-day-old Col-0 wild type and *hda15-1* were treated with or without 50 μM ABA for 3 hr. Total proteins were extracted in an extraction buffer (50 mM Tris-HCl, pH 7.4, 150 mM NaCl, 10% glycerol, 1% Igepal CA-630, and 1 mM PMSF) containing a protease inhibitor cocktail (Roche). The HDA15-associtated proteins were immunoprecipitated by rabbit polyclonal anti-HDA15 (Liu *et al.*, 2013). After trypsin digestion and desalting, peptides were used to perform LC-MS/MS by Thermo Orbitrap Elite Mass Spectrometer and for Mascot analysis. *hda15-1* was used as a negative control for excluding the proteins that bind non-specifically to the HDA15 antibody in immunopurification. The correlation of the interacting proteins was analyzed by STRING (Szklarczyk et al., 2015) and visualized in Cytoscape (Shannon et al., 2003).

### Bimolecular fluorescence complementation assays

HDA15, MAC3A, and MAC3B were cloned into the pCR8/GW/TOPO vector and then recombined into the YFP^N^ (pEarleyGate201-YFP^N^) and YFP^C^ (pEarleyGate202-YFP^C^) vectors (Lu et al., 2010). Constructed vectors were transiently transformed into Arabidopsis protoplasts by polyethylene-glycol (PEG)-mediated transfection (Yoo *et al.*, 2007). YFP florescence was observed using Leica TCS SP5 confocal microscope.

### Co-immunoprecipitation assays

MAC3A or MAC3B cDNA was cloned into the pK7WGF2 binary vector for transient expressing GFP-MAC3A or GFP-MAC3B by PEG-mediated transfection in Col-0 Arabidopsis protoplasts. 10-day-old Col-0, *hda15-1*, and *Pro35S:GFP-MAC3B* transgenic plants were treated with or without 50 μM ABA for 3 hr. Total proteins were extracted in the extraction buffer containing protease inhibitor cocktail (Roche) as described above. Protein extracts were incubated with a rabbit polyclonal anti-HDA15 antibody and protein G Mag Sepharose beads (GE Healthcare) overnight at 4°C. The beads were washed with a wash buffer (50 mM Tris-HCl, pH 7.4, 150 mM NaCl, 10% glycerol, and 1% Igepal CA-630). Immunoblotting was carried out by using the rabbit polyclonal anti-HDA15 antibody, anti-GFP antibody (Abcam, ab290), and the secondary antibody Clean-Blot IP Detection Reagent (Thermo, 21230).

### Abiotic stress tolerance test and seed germination treated with ABA

For the salt tolerance test, 3-week-old plants were watered with 300 mM NaCl for two weeks, and the survival rates were measured. For transpiration (water loss) measurements, detached leaves from 5-week-old plants were exposed to white light in a growth chamber at 23°C. Leaves were weighed at various time intervals, and the loss of fresh weight (percentage) was used to indicate water loss. For seed germination test, imbibed seeds were cold treated at 4°C in the dark for 3 days, and then sown on a half-strength Murashige and Skoog (MS) medium (pH5.7) with 1% sucrose, 0.8% w/v agar, and 2 μM ABA for 3 days.

### RNA-seq

For RNA sequencing, 10-day-old seedlings of Col-0 wild type, *hda15-1*, and *mac3a/mac3b* were treated with or without 50 μM ABA for 3 hr. Total RNA was extracted with Trizol reagent (Invitrogen) according to the manufacturer’s protocol, and then used for Illumina NovaSeq 6000 platform to generate paired-end reads of 150 bp. Two biological replicates for each sample were performed independently.

After low quality read trimming, over 73 million paired-end reads were obtained per sample. More than 94% of the reads for every replicate were mapped to Araport11 using Bowtie 2 (Langmead & Salzberg, 2012) and BLAT (Kent, 2002). RPKM values were calculated using the RackJ software package (http://rackj.sourceforge.net/), and normalized by the TMM method (Robinson & Oshlack, 2010). In this study, differentially expressed genes were defined as *p*-value <0.05 (*t*-test) with fold-change≥ 1.5.

The major types of AS events, IR, AltD/A, and ES, were analyzed by RackJ as described previously (Kanno *et al.*, 2017). In brief, significant IR changes were defined as *p*-value<0.05 (*t*-test) with fold-change> 2. The significant ES and AltD/A events were determined using a method similar to that for IR events following the criteria: the read counts supporting the ES or AltD/A event >20, the read counts aligned to the skipped exon >20 (for ES); the read counts supporting all other junctions of the same intron (for AltD/A).

### qRT-PCR

Total RNA was extracted with Trizol reagent (Invitrogen) according to the manufacturer’s protocol and used to synthesize cDNA. qRT-PCR was performed using iQ SYBR Green Supermix (Bio-Rad) and the CFX96 real-time PCR system (Bio-Rad). The gene-specific primers used for real-time PCR are listed in Supplemental Data Set 9. Each sample was quantified at least in triplicate and normalized using *Ubiquitin10* (*UBQ10*) as an internal control.

### ChIP-qPCR

ChIP assays were performed as described (Gendrel *et al.*, 2005; Liu *et al.*, 2013). Chromatin was extracted from 10-day-old seedlings. After fixation with 1% formaldehyde, the chromatin was sheared to an average length of 500 bp by sonication and then immunoprecipitated with the H3K9ac antibody (Diagenode, C15410004). The cross-linking was then reversed, and the amount of each precipitated DNA fragment was determined by qPCR using specific primers in Supplemental Data set 9. Three biological replicates were performed, and three technical repeats were carried out for each biological replicate. Representative results from one biological replicate were shown.

### Accession numbers

HDA15, AT3G18520; MAC3A, AT1G04510; MAC3B, AT2G33340; SNRK2.6, AT4G33950; NTAQ1, AT2G41760; EDM1, AT4G11260; UBQ10, AT4G05320. All RNA-seq data have been uploaded to the Sequence Read Archive (SRA) database (accession no. SRP292622) at the National Center for Biotechnology Information (NCBI).

## Supplemental Data Files

**Supplemental Figure 1** Biological processes of HDA15 interacting proteins in mock and ABA treatment

**Supplemental Data Set 1.** HDA15 interacting proteins in mock and ABA treatment identified by LC-MS/MS

**Supplemental Data Set 2.** Differentially expressed genes in *hda15-1* compared to Col-0 in mock and ABA treatment.

**Supplemental Data Set 3.** Differentially expressed genes in *mac3a/mac3b* compared to Col-0 in mock and ABA treatment.

**Supplemental Data Set 4.** Coregulated genes in *hda15-1* and *mac3a/mac3b* in mock and ABA treatment.

**Supplemental Data Set 5.** List of IR events in *hda15-1* and *mac3a/mac3b* in mock and ABA-treated conditions compared with Col-0.

**Supplemental Data Set 6.** List of ES events in *hda15-1* and *mac3a/mac3b* in mock and ABA-treated conditions compared with Col-0.

**Supplemental Data Set 7.** List of AltD/A events in *hda15-1* and *mac3a/mac3b* in mock and ABA-treated conditions compared with Col-0.

**Supplemental Data Set 8.** List of AS events in Col-0, *hda15-1* and *mac3a/mac3b* in ABA-treated condition compared with mock conditions.

**Supplemental Data Set 9.** List of primers used in this study.

## Acknowledgments

We are grateful to the Technology Commons, College of Life Science, National Taiwan University for the convenient use of the Bio-Rad real-time PCR system and Leica TCS SP5 confocal microscope. We also thank the Bioinformatics Core Lab of the Institute of Plant and Microbial Biology at Academia Sinica, for the data analysis of alternative splicing. This work was supported by the Ministry of Science and Technology of the Republic of China (108-2311-B-002-013-MY3 and 109-2311-B-002-027 to K. W., and 109-2313-B-001 −009 −MY3 to P.-Y.C) and NTU-Academia Sinica joint grant (NTU-AS-108L104310 to K.W. and P.-Y.C).

## Author contributions

Y.-T. T., P.-Y. C and K. W. designed the study. Y.-T. T., C.-Y. C., Y.-S. H., M.-R. Y. and J.-W. A. H. performed the experiments and analyzed the data. Y.-T. T., P.-Y. C and K. W. wrote and edited the manuscript.

**Supplemental Fig. 1.**
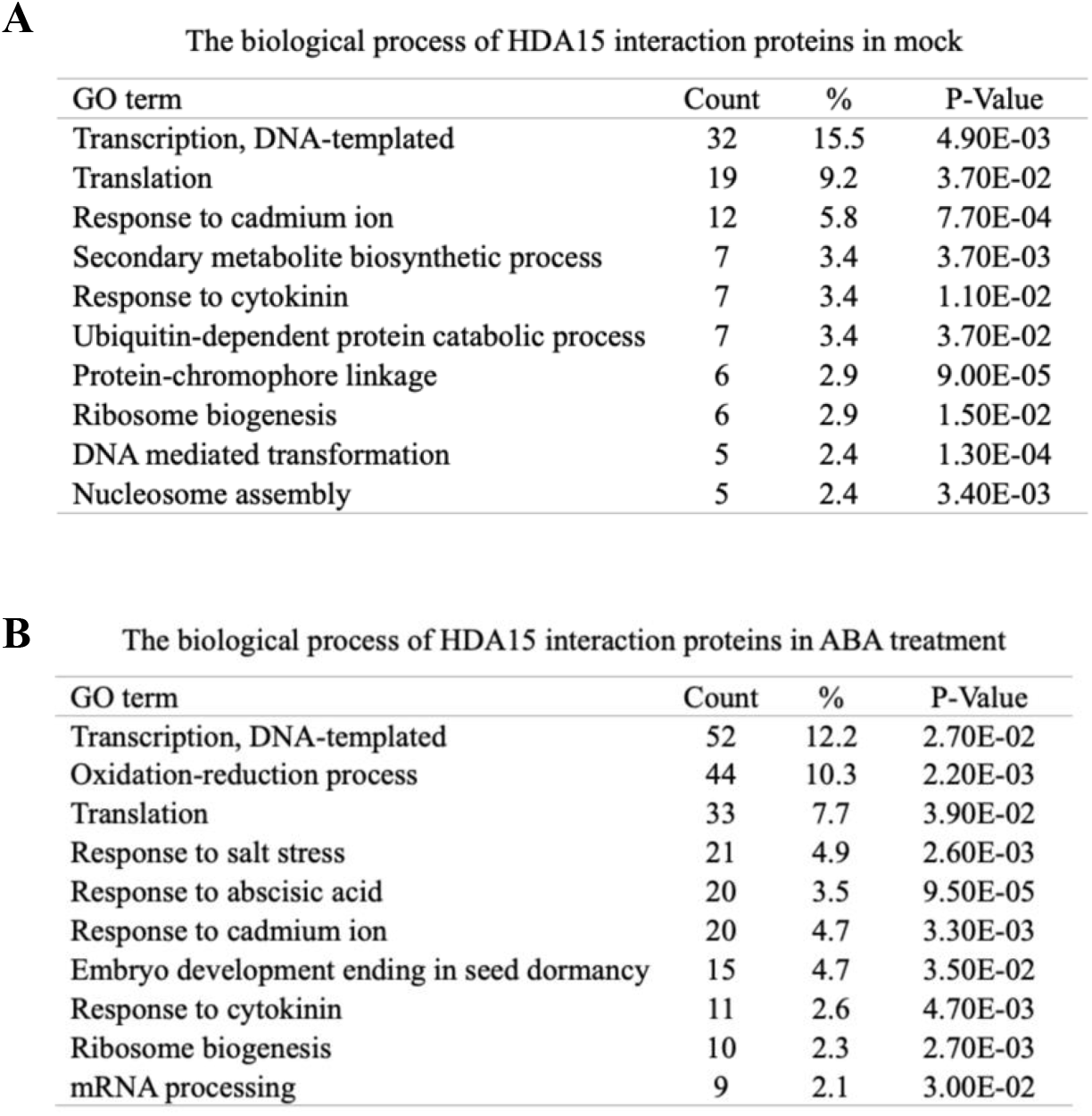
Biological processes of HDA15 interacting proteins in mock and ABA treatment

## Parsed Citations

Abe, H., Urao, T., Ito, T., Seki, M., Shinozaki, K., and Yamaguchi-Shinozaki, K. (2003). Arabidopsis AtMYC2 (bHLH) and AtMYB2 (MYB) function as transcriptional activators in abscisic acid signaling. The Plant cell 15, 63–78. Google Scholar: Author Only Title Only Author and Title

Caro, E., Stroud, H., Greenberg, M.V., Bernatavichute, Y.V., Feng, S., Groth, M., Vashisht, A.A., Wohlschlegel, J., and Jacobsen, S.E. (2012). The SET-domain protein SUVR5 mediates H3K9me2 deposition and silencing at stimulus response genes in a DNA methylation-independent manner. PLoS genetics 8, e1002995. Google Scholar: Author Only Title Only Author and Title

Chan, S.P., Kao, D.I., Tsai, W.Y., and Cheng, S.C. (2003). The Prp19p-associated complex in spliceosome activation. Science 302, 279–282. Google Scholar: Author Only Title Only Author and Title

Chen, C.Y., Tu, Y.T., Hsu, J.C., Hung, H.C., Liu, T.C., Lee, Y.H., Chou, C.C., Cheng, Y.S., and Wu, K. (2020). Structure of Arabidopsis HISTONE DEACETYLASE15. Plant physiology. Google Scholar: Author Only Title Only Author and Title

Chen, Z.J., and Tian, L. (2007). Roles of dynamic and reversible histone acetylation in plant development and polyploidy. Biochimica et biophysica acta 1769, 295–307. Google Scholar: Author Only Title Only Author and Title

Fisher, A.J., and Franklin, K.A. (2011). Chromatin remodelling in plant light signalling. Physiologia plantarum 142, 305–313. Google Scholar: Author Only Title Only Author and Title

Gu, D., Chen, C.Y., Zhao, M., Zhao, L., Duan, X., Duan, J., Wu, K., and Liu, X. (2017). Identification of HDA15-PIF1 as a key repression module directing the transcriptional network of seed germination in the dark. Nucleic acids research 45, 7137–7150. Google Scholar: Author Only Title Only Author and Title

Gunderson, F.Q., and Johnson, T.L. (2009). Acetylation by the transcriptional coactivator Gcn5 plays a novel role in co-transcriptional spliceosome assembly. PLoS genetics 5, e1000682. Google Scholar: Author Only Title Only Author and Title

Gunderson, F.Q., Merkhofer, E.C., and Johnson, T.L. (2011). Dynamic histone acetylation is critical for cotranscriptional spliceosome assembly and spliceosomal rearrangements. Proceedings of the National Academy of Sciences of the United States of America 108, 2004–2009. Google Scholar: Author Only Title Only Author and Title

Hogg, R., McGrail, J.C., and O’Keefe, R.T. (2010). The function of the NineTeen Complex (NTC) in regulating spliceosome conformations and fidelity during pre-mRNA splicing. Biochem Soc Trans 38, 1110–1115. Google Scholar: Author Only Title Only Author and Title

Jia, T., Zhang, B., You, C., Zhang, Y., Zeng, L., Li, S., Johnson, K.C.M., Yu, B., Li, X., and Chen, X. (2017). The Arabidopsis MOS4-Associated Complex Promotes MicroRNABiogenesis and Precursor Messenger RNASplicing. The Plant cell 29, 2626–2643. Google Scholar: Author Only Title Only Author and Title

Kim, J.M., Sasaki, T., Ueda, M., Sako, K., and Seki, M. (2015). Chromatin changes in response to drought, salinity, heat, and cold stresses in plants. Frontiers in plant science 6, 114. Google Scholar: Author Only Title Only Author and Title

Lee, H.G., and Seo, P.J. (2019). MYB96 recruits the HDA15 protein to suppress negative regulators of ABA signaling in Arabidopsis. Nature Communications 10. Google Scholar: Author Only Title Only Author and Title

Li, S., Liu, K., Zhou, B., Li, M., Zhang, S., Zeng, L., Zhang, C., and Yu, B. (2018). MAC3A and MAC3B, Two Core Subunits of the MOS4-Associated Complex, Positively Influence miRNA Biogenesis. The Plant cell 30, 481–494. Google Scholar: Author Only Title Only Author and Title

Li, Y., Yang, J., Shang, X., Lv, W., Xia, C., Wang, C., Feng, J., Cao, Y., He, H., Li, L., and Ma, L. (2019). SKIP regulates environmental fitness and floral transition by forming two distinct complexes in Arabidopsis. The New phytologist 224, 321–335. Google Scholar: Author Only Title Only Author and Title

Liu, X., Yang, S., Zhao, M., Luo, M., Yu, C.W., Chen, C.Y., Tai, R., and Wu, K. (2014). Transcriptional repression by histone deacetylases in plants. Molecular plant 7, 764–772. Google Scholar: Author Only Title Only Author and Title

Liu, X., Chen, C.Y., Wang, K.C., Luo, M., Tai, R., Yuan, L., Zhao, M., Yang, S., Tian, G., Cui, Y., Hsieh, H.L., and Wu, K. (2013). PHYTOCHROME INTERACTING FACTOR3 associates with the histone deacetylase HDA15 in repression of chlorophyll biosynthesis and photosynthesis in etiolated Arabidopsis seedlings. The Plant cell 25, 1258–1273. Google Scholar: Author Only Title Only Author and Title

Liu, Y., Zhang, A., Yin, H., Meng, Q., Yu, X., Huang, S., Wang, J., Ahmad, R., Liu, B., and Xu, Z.Y. (2018). Trithorax-group proteins ARABIDOPSIS TRITHORAX4 (ATX4) and ATX5 function in abscisic acid and dehydration stress responses. The New phytologist 217, 1582–1597. Google Scholar: Author Only Title Only Author and Title

Luco, R.F., Pan, Q., Tominaga, K., Blencowe, B.J., Pereira-Smith, O.M., and Misteli, T. (2010). Regulation of alternative splicing by histone modifications. Science 327, 996–1000. Google Scholar: Author Only Title Only Author and Title

Mehdi, S., Derkacheva, M., Ramstrom, M., Kralemann, L., Bergquist, J., and Hennig, L. (2016). The WD40 Domain Protein MSI1 Functions in a Histone Deacetylase Complex to Fine-Tune Abscisic Acid Signaling. The Plant cell 28, 42–54. Google Scholar: Author Only Title Only Author and Title

Monaghan, J., Xu, F., Xu, S., Zhang, Y., and Li, X. (2010). Two putative RNA-binding proteins function with unequal genetic redundancy in the MOS4-associated complex. Plant physiology 154, 1783–1793. Google Scholar: Author Only Title Only Author and Title

Monaghan, J., Xu, F., Gao, M., Zhao, Q., Palma, K., Long, C., Chen, S., Zhang, Y., and Li, X. (2009). Two Prp19-like U-box proteins in the MOS4-associated complex play redundant roles in plant innate immunity. PLoS pathogens 5, e1000526. Google Scholar: Author Only Title Only Author and Title

Pajoro, A., Severing, E., Angenent, G.C., and Immink, R.G.H. (2017). Histone H3 lysine 36 methylation affects temperature-induced alternative splicing and flowering in plants. Genome Biol 18, 102. Google Scholar: Author Only Title Only Author and Title

Palma, K., Zhao, Q., Cheng, Y.T., Bi, D., Monaghan, J., Cheng, W., Zhang, Y., and Li, X. (2007). Regulation of plant innate immunity by three proteins in a complex conserved across the plant and animal kingdoms. Genes & development 21, 1484–1493. Google Scholar: Author Only Title Only Author and Title

Pandey, R., Müller, A., Napoli, C.A., Selinger, D.A., Pikaard, C.S., Richards, E.J., Bender, J., Mount, D.W., and Jorgensen, R.A. (2002). Analysis of histone acetyltransferase and histone deacetylase families of Arabidopsis thaliana suggests functional diversification of chromatin modification among multicellular eukaryotes. Nucleic acids research 30, 5036–5055. Google Scholar: Author Only Title Only Author and Title

Piacentini, L., Fanti, L., Negri, R., Del Vescovo, V., Fatica, A., Altieri, F., and Pimpinelli, S. (2009). Heterochromatin protein 1 (HP1a) positively regulates euchromatic gene expression through RNAtranscript association and interaction with hnRNPs in Drosophila. PLoS genetics 5, e1000670. Google Scholar: Author Only Title Only Author and Title

Rahhal, R., and Seto, E. (2019). Emerging roles of histone modifications and HDACs in RNAsplicing. Nucleic acids research 47, 4911–4926. Google Scholar: Author Only Title Only Author and Title

Ren, X., Chen, Z., Liu, Y., Zhang, H., Zhang, M., Liu, Q., Hong, X., Zhu, J.K., and Gong, Z. (2010). ABO3, a WRKY transcription factor, mediates plant responses to abscisic acid and drought tolerance in Arabidopsis. The Plant journal : for cell and molecular biology 63, 417–429. Google Scholar: Author Only Title Only Author and Title

Ruhl, C., Stauffer, E., Kahles, A., Wagner, G., Drechsel, G., Ratsch, G., and Wachter, A. (2012). Polypyrimidine tract binding protein homologs from Arabidopsis are key regulators of alternative splicing with implications in fundamental developmental processes. The Plant cell 24, 4360–4375. Google Scholar: Author Only Title Only Author and Title

Saint-André, V., Batsché, E., Rachez, C., and Muchardt, C. (2011). Histone H3 lysine 9 trimethylation and HP1γ favor inclusion of alternative exons. Nature structural & molecular biology 18, 337–344. Google Scholar: Author Only Title Only Author and Title

Shannon, P., Markiel, A., Ozier, O., Baliga, N.S., Wang, J.T., Ramage, D., Amin, N., Schwikowski, B., and Ideker, T. (2003). Cytoscape: a software environment for integrated models of biomolecular interaction networks. Genome research 13, 2498–2504. Google Scholar: Author Only Title Only Author and Title

Shen, Y., Lei, T., Cui, X., Liu, X., Zhou, S., Zheng, Y., Guérard, F., Issakidis-Bourguet, E., and Zhou, D.X. (2019). Arabidopsis histone deacetylase HDA15 directly represses plant response to elevated ambient temperature. The Plant journal : for cell and molecular biology 100, 991–1006. Google Scholar: Author Only Title Only Author and Title

Sims, R.J., 3rd, Millhouse, S., Chen, C.F., Lewis, B.A., Erdjument-Bromage, H., Tempst, P., Manley, J.L., and Reinberg, D. (2007). Recognition of trimethylated histone H3 lysine 4 facilitates the recruitment of transcription postinitiation factors and pre-mRNA splicing. Molecular cell 28, 665–676. Google Scholar: Author Only Title Only Author and Title

Szklarczyk, D., Franceschini, A., Wyder, S., Forslund, K., Heller, D., Huerta-Cepas, J., Simonovic, M., Roth, A., Santos, A., Tsafou, K.P., Kuhn, M., Bork, P., Jensen, L.J., and von Mering, C. (2015). STRING v10: protein-protein interaction networks, integrated over the tree of life. Nucleic acids research 43, D447–452. Google Scholar: Author Only Title Only Author and Title

Tang, Y., Liu, X., Liu, X., Li, Y., Wu, K., and Hou, X. (2017). Arabidopsis NF-YCs Mediate the Light-Controlled Hypocotyl Elongation via Modulating Histone Acetylation. Molecular plant 10, 260–273. Google Scholar: Author Only Title Only Author and Title

Ueda, M., and Seki, M. (2020). Histone Modifications Form Epigenetic Regulatory Networks to Regulate Abiotic Stress Response. Plant physiology 182, 15–26. Google Scholar: Author Only Title Only Author and Title

Wang, Z., Ji, H., Yuan, B., Wang, S., Su, C., Yao, B., Zhao, H., and Li, X. (2015). ABA signalling is fine-tuned by antagonistic HAB1 variants. Nat Commun 6, 8138. Google Scholar: Author Only Title Only Author and Title

Xu, Y., Gan, E.S., Zhou, J., Wee, W.Y., Zhang, X., and Ito, T. (2014). Arabidopsis MRG domain proteins bridge two histone modifications to elevate expression of flowering genes. Nucleic acids research 42, 10960–10974. Google Scholar: Author Only Title Only Author and Title

Yu, C.W., Tai, R., Wang, S.C., Yang, P., Luo, M., Yang, S., Cheng, K., Wang, W.C., Cheng, Y.S., and Wu, K. (2017). HISTONE DEACETYLASE6 Acts in Concert with Histone Methyltransferases SUVH4, SUVH5, and SUVH6 to Regulate Transposon Silencing. The Plant cell 29, 1970–1983. Google Scholar: Author Only Title Only Author and Title

Zhang, S., Liu, Y., and Yu, B. (2014). PRL1, an RNA-binding protein, positively regulates the accumulation of miRNAs and siRNAs in Arabidopsis. PLoS genetics 10, e1004841. Google Scholar: Author Only Title Only Author and Title

Zhao, L., Peng, T., Chen, C.Y., Ji, R., Gu, D., Li, T., Zhang, D., Tu, Y.T., Wu, K., and Liu, X. (2019). HY5 Interacts with the Histone Deacetylase HDA15 to Repress Hypocotyl Cell Elongation in Photomorphogenesis. Plant physiology 180, 1450–1466. Google Scholar: Author Only Title Only Author and Title

Zhu, J., Jeong, J.C., Zhu, Y., Sokolchik, I., Miyazaki, S., Zhu, J.K., Hasegawa, P.M., Bohnert, H.J., Shi, H., Yun, D.J., and Bressan, R.A. (2008). Involvement of Arabidopsis HOS15 in histone deacetylation and cold tolerance. Proceedings of the National Academy of Sciences of the United States of America 105, 4945–4950. Google Scholar: Author Only Title Only Author and Title

